# Lgr5^+^ ductal cells of von Ebner’s glands are stem cells for turnover of posterior tongue taste buds

**DOI:** 10.1101/2024.04.18.589316

**Authors:** Theresa A. Harrison, Anthony M. Downs, Alexandria J. Slepian, Johan H. van Es, Hans Clevers, Dennis M. Defoe

**Affiliations:** Department of Biomedical Sciences, Quillen College of Medicine, East Tennessee State University, Johnson City, TN, United States of America; Hubrecht Institute, Royal Netherlands Academy of Arts and Sciences (KNAW) and University Medical Center (UMC) Utrecht, Uppsalalaan 8, 3584 CT Utrecht, the Netherlands

**Keywords:** Circumvallate, foliate, Cre, Lgr5, mTmG, salivary gland ducts, taste bud, taste papilla, von Ebner’s, Wnt

## Abstract

Taste bud cells have a limited lifespan and must be continuously replaced along with the papilla epithelium in which they reside. Previous work has shown that expression of leucine-rich G protein-coupled receptor 5 (Lgr5), a Wnt pathway agonist, serves as a marker of adult stem/progenitor cells for taste buds located in posterior tongue (circumvallate and foliate), but not anterior tongue (fungiform), taste papillae. However, the specific location/niche of the *Lgr5*-expressing cells supporting renewal and their phenotypic properties have not been fully explored. To address this, the genesis and fate of Lgr5^+^ cells were examined in developing and adult mice using genetic reporter strains. Evidence from *Lgr5*-*lacZ* and *Lgr5*- *GFP* mice shows that, while *Lgr5* is broadly expressed in the epithelium of nascent circumvallate papillae and their trenches during embryonic development, it becomes concentrated within the ducts of adjacent von Ebner’s salivary glands during the first postnatal week, co-incident with the appearance of differentiated taste buds. In posterior tongue taste papillae of adult animals, sites of highest *Lgr5* expression are found in excretory ducts, restricted to the outer (basal) layer of the bi-layered excretory zone. These Lgr5^+^ cells are immunoreactive for keratin 14, like cells in the basal layer of extragemmal taste epithelium, and for Sox9, a marker of exocrine gland duct cells. Lineage tracing experiments with an *Lgr5*-*EGFP*-*IRES*-*CreERT2;R26*-*mTmG* reporter show that Lgr5^+^ ductal cells become labeled one day following Cre induction, prior to the appearance of descendent cells in taste buds and extragemmal epithelium. These data support a role for Lgr5^+^ ductal cells as stem cells and suggest that a cooperative interaction exists between posterior taste epithelium and its associated salivary glands in taste cell turnover.

## Introduction

Mammalian taste buds are complex sensory organs consisting of taste receptor cells that interact with each other via local neurochemical circuits and ultimately communicate with sensory nerves via both conventional synaptic and non-synaptic interactions (Finger et al., 2005; Ma et al., 2018; Romanov et al., 2018). Along with supporting cells and immature precursors, these receptor cells are arranged as compact clusters (taste buds) embedded within the stratified surface epithelium of the tongue, palate and epiglottis. Lingual taste buds, along with surrounding epithelium and connective tissue, are organized into structures called papillae that are distributed non-uniformly over the dorsal tongue surface. Mouse tongues have three major regions that contain taste papillae: the anterior tongue, across which fungiform papillae are distributed in a patterned array, two sets of foliate papillae located on opposite lateral edges of the posterior tongue, and a single circumvallate papilla that lies in the midline near the back of the tongue.

While taste buds in different tongue regions are structurally very similar, their arrangement within papillae differs markedly. Fungiform papillae are relatively simple dome-shaped structures consisting of a surface epithelium, each with a single taste bud, that envelopes a connective tissue core. By contrast, in circumvallate and foliate papillae several hundred buds occupy the walls of trenches, infoldings formed by epithelial invaginations from the tongue surface. Posterior papillae are also distinguished from anterior papillae by their connection to serous salivary glands, the von Ebner’s glands, which are located deeper in the tongue. These glands empty their contents into the luminal spaces of papilla trenches via excretory ducts. In addition to their structural differences, anterior and posterior papillae have distinct embryological origins. While fungiform papillae are presumed to arise from ectoderm, both circumvallate and foliate papillae are endodermal derivatives (Rothova et al., 2012) that develop in tandem with their associated salivary glands (Jitpukdeebodintra et al., 2002; Lee et al., 2006; Sbarbati et al., 2000,2001; Sohn et al., 2011).

The peripheral taste system of vertebrates is unusual among sensory pathways in that its cellular components are continually turned over in adult animals (Beidler and Smallman, 1965; Conger and Wells, 1969; Farbman, 1980). The molecular signaling mechanisms regulating homeostasis in adults include several that are shared with taste bud development (Barlow, 2015; Krimm and Barlow, 2015). Prominent among these is the canonical Wnt pathway, which plays a key role in both formation and maintenance of taste papillae (Liu et al., 2006; Iwatsuki et al., 2007; Gaillard et al., 2017). Wnt-driven transcription, initiated through interaction of nuclear β-catenin with Tcf/Lef transcriptional cofactors, is part of an integrated program for renewal and regeneration in many tissues (Clevers et al., 2014). One of the targets under Wnt control, *leucine-rich G protein-coupled receptor 5* (*Lgr5*), has been shown to be a marker for stem cells in several different areas of the body (Barker et al., 2007, 2010; Jaks et al., 2008; Plaks et al., 2013; Rios et al., 2014), including the posterior tongue (Takeda et al., 2014; Yee et al., 2014). In lineage tracing experiments, Lgr5^+^ cells were shown to be responsible for generation of all taste cell types during normal homeostasis and for replacement of taste buds following neurectomy-induced degeneration (Takeda et al., 2014; Yee et al., 2014). In these studies, however, characterization of the long-term progenitors giving rise to taste cells and the niche in which they reside was not undertaken.

In the present report, we have examined *Lgr5* expression and lineage of Lgr5^+^ cells in the tongue of developing and adult mice. Our studies establish excretory ducts of von Ebner’s glands, associated with circumvallate and foliate papillae, as the major lingual sites of *Lgr5* expression in adult animals. Found only in the outer layer of the double-layered excretory ducts and situated at their confluence with papilla epithelium, these Lgr5^+^ cells are unique in exhibiting a set of phenotypic marker proteins that are typically differentially expressed between ductal and papillary epithelial tissues. Lineage analysis shows that ductal Lgr5^+^ cells undergo long-term renewal and give rise to differentiated cells of both taste buds and of their surrounding stratified epithelium; thus, they are likely to represent the primary stem/progenitor cells for posterior taste buds of adults.

## Materials and Methods

### Ethics statement

Experimental procedures involving animals were approved by the institutional review committees at East Tennessee State University (ETSU) and the Hubrecht Institute. They also complied with the National Institutes of Health Guidelines for Care and Use of Animals in Research and with the recommendations of ARRIVE (Animal Research: Reporting of Experiments; https://arriveguidelines.org).

### Mice

*Lgr5*-*IRES*-*lacZ* mice were resident at the Hubrecht Institute. These animals possess an *IRES*-*lacZ* cassette targeted to the 5′ end of the last exon, removing all transmembrane regions of the encoded Lgr5 protein and placing the β-galactosidase enzyme (lacZ) under the *Lgr5* control region (Barker et al., 2007). Non-transgenic mice (C57Bl/J6), as well as *Lgr5*-*EGFP*-*IRES*-*CreERT2* and *Rosa26*-*td*-*Tomato,−EGFP* (*mTmG*) strains, were obtained from the Jackson Labs (Bar Harbor, ME) and housed and bred at facilities in the Division of Laboratory Animal Resources at ETSU. *Lgr5*-*EGFP*-*IRES*-*CreERT2* mice carry a knock-in allele that both abolishes *Lgr5* gene function and expresses EGFP and CreERT2 fusion proteins from *Lgr5* promoter/enhancer elements. The *mTmG* line, a two-color fluorescent Cre-reporter, possesses *loxP* sites on either side of an expression cassette (inserted into the ROSA 26 locus) targeting tdTomato (mT) to cell membranes, causing strong red fluorescence labeling in all tissues and cell types. When bred to mice with a tamoxifen- activable Cre recombinase, the resulting offspring have the *mT* cassette deleted in activated cells and the lineages derived from them, allowing expression of membrane-targeted EGFP (mG) via an expression cassette located immediately downstream. Maintenance and genotyping of mice has been described previously (Barker et al., 2007; Muzumdar et al., 2007).

For most expression studies, the *Lgr5*-*EGFP*-*IRES*-*CreERT2* and *Lgr5*-*IRES*-*lacZ* lines were bred to C57BL/6J mice to produce heterozygotes (referred to hereafter as *Lgr5*-*GFP* and *Lgr5*-*lacZ*). In the case of lineage tracing, *Lgr5*-*EGFP*-*IRES*-*CreERT2* heterozygotes were crossed with homozygous *Rosa26*-*td*-*Tomato,−EGFP* (*mTmG*) animals and progeny heterozygous for both transgenes were selected for experiments. Uninduced animals with the same genotype (*Lgr5^EGFP^*^-*IRES*-*CreERT*2/+;*R26R*-*mTmG/+*^) were also used in some *Lgr5* expression experiments.

To initiate Cre induction, mice were injected intraperitoneally with a single 200 µl dose of tamoxifen (20 mg ml^-1^; Sigma Chemical Co., St. Louis, MO) in corn oil (Madisen et al., 2010). Control animals received corn oil alone.

### Tissue preparation and histological staining

Mice were euthanized by CO_2_ asphyxiation and tongues surgically removed. For fluorescent protein and immunocytochemical visualization, whole tongues were immersion-fixed overnight at 4°in 2% paraformaldehyde in 0.1 M sodium acetate buffer, pH 6.0 (Lorincz and Nusser, 2010). For lacZ staining, fixation involved immersion at 4°in 4% paraformaldehyde in phosphate-buffered saline, pH 7.3 (PBS). Following brief rinsing in PBS, the tongues were either left intact for documentation of GFP fluorescence in whole-mounts (see below) or further dissected into anterior, posterior and lateral pieces, containing fungiform, circumvallate and foliate papillae, respectively, and prepared for sectioning (see below).

For experiments on isolated circumvallate papillae, the extirpated tongues from non-transgenic and *Lgr5*-*EGFP*-*IRES*-*CreERT2;R26*-*mTmG* reporter mice were injected with approximately 100-200 μl 1% Dispase II (neutral protease; Roche, Indianapolis, IN) dissolved in Balanced Salt Solution (BSS) (Ozdener et al., 2006; Harrison et al., 2011). Injections were localized to the intradermal space immediately beneath the papilla. Following incubation for 15 minutes at room temperature in BSS, the epidermal layer, including that of the papilla, was peeled away and extraneous tissue trimmed. The papilla epidermis was then fixed at room temperature in 4% paraformaldehyde in PBS, rinsed and examined by fluorescence stereomicroscopy.

Paraffin embedding and sectioning, as well tissue staining for the presence of lacZ activity, have been described (Barker et al., 2007). Processing of dissected tongue specimens for frozen sectioning and procedures used for immunolabeling of sections followed previously published protocols (Harrison et al., 2016). Primary antibodies used included rabbit anti-cytokeratin 14 (Thermo Scientific, Waltham, MA; 1:250), rat anti-cytokeratin 8 (TROMA-I; Developmental Studies Hybridoma Bank; 1:500) and rabbit anti- Sox9 (Millipore, Burlington, MA; 1:750). Detection of antibody binding was with the appropriate species-specific Biotin-SP-conjugated goat anti-antibodies (1:400) followed by AlexaFluor 555-conjugated Streptavidin (1:600) (Life Technologies, Carlsbad, CA; 1:400).

All studies of *Lgr5* expression, through its surrogate reporters, exploited the strong lacZ and GFP signals from the *Lgr5*-*IRES-lacZ* and *Lgr5*-*EGFP*-*IRES-CreERT2* transgenes. No labeled antibodies were used for intensification of the expressed enzyme and fluorescent proteins.

### Microscopy

LacZ reaction product in 5 μm paraffin sections was viewed on an Olympus BH-2 light microscope (Olympus, Center Valley, PA) and images captured and digitized using an Insight CCT camera (Model 142 Color Mosaic) and Spot Software System (Diagnostic Instruments, Sterling Heights, MI). Fluorescence images of whole-mounted tongues and isolated papillae were obtained using a Leica MZ16FA stereomicroscope with a Fluo III fluorescence module (Leica, Heidelberg, Germany) and documented with a color digital CCD camera and image capture software (Q Imaging Retiga EXi camera and Q Capture software, Surrey, Canada). Frozen sections (10 μm) were examined in Leica SP2 and SP8 laser scanning confocal microscopes (Leica, Heidelberg, Germany) using 20X and 60X infinity-corrected objectives and viewed through the GFP and RFP fluorescence channels or using manufacturer-provided Alexa 488 and Alexa 555 excitation and emission settings, as appropriate for the labeling paradigm used. The resulting images were processed using Photoshop CS2 (Adobe Systems, San Jose, CA).

## Results

### *Lgr5* expression by developing taste papillae

*Lgr5* expression was initially examined in tongue whole-mounts from neonatal and adult mice with an *Lgr5*-*EGFP*-*IRES*-*CreERT2* allele (*Lgr5*-*GFP* reporter mice). Consistent with a previous report (Takeda et al., 2013), in these mice robust GFP labeling demarcates all three types of lingual taste papillae at 1 week after birth, when taste buds have begun forming at all sites (Supplemental Fig. 1A, B; Krimm et al., 2015). Expression in circumvallate and foliate papillae is maintained at high levels in 12 week-old animals (Supplemental Fig. 1C), but is undetectable in fungiform papillae at this age, either in tongue whole-mounts (Supplemental Fig. 1D) or histological sections (data not shown; Takeda et al., 2013).

To characterize the distribution of Lgr5^+^ cells in developing circumvallate papillae, sections of posterior tongue from *Lgr5*-*GFP* reporter animals were viewed at selected times from mid-gestation to the initial postnatal period. Early in papilla morphogenesis, *Lgr5* expression in the central posterior tongue is restricted to columnar cells of the presumptive taste epithelium, which forms a continuous single layer that covers the surface of the dome-like central portion and lines the invaginations of the developing papilla trenches (Fig. 1A). By late gestation (Fig. 1B), labeling of the multilayered epithelium appears patchy due to an irregular distribution of Lgr5^+^ cells. At this stage, Lgr5^+^ cells are found more deeply in the mesenchyme, both in the lengthening invaginations of the papilla and in the forming ducts of von Ebner’s salivary glands that extend inferior to the papilla itself (Fig. 1B). At birth (Fig. 1C, D), expression is still evident in the medial and lateral walls of the trenches and in the epithelium of nascent ducts. However, by 1 week postnatally (Fig. 1E, F), conspicuous concentrations of GFP fluorescence appear below the trenches, in regions where salivary gland ducts meet the wall of the taste papilla. By this stage, these are the areas of highest *Lgr5* expression associated with the papilla.

**Figure 1.**
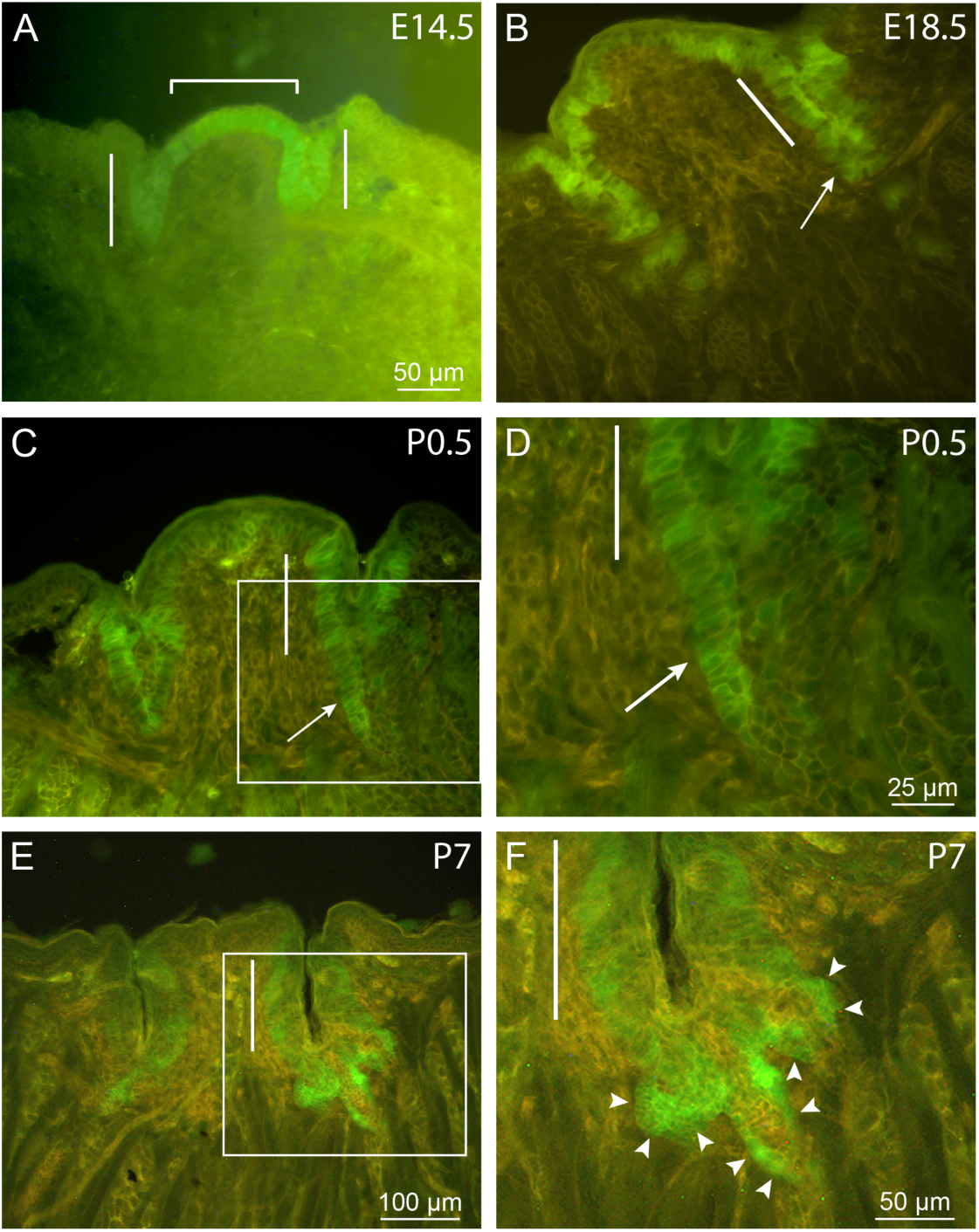
*Lgr5* expression by developing circumvallate papillae of *Lgr5*-*GFP* knock-in mice. (A) E14.5. A layer of GFP-labeled epithelial cells is observed covering the papilla dome (bracket) and lining the invaginations of the nascent trenches (bars). (B) E18.5. Labeling within the papilla epithelium is interrupted by non-expressing cells in both the dome and trench areas. Lgr5^+^ cells are also found deeper in the mesenchyme, associated with developing salivary ducts (arrow). At birth (C), GFP fluorescence is still evident in the medial and lateral walls of the papilla trenches (e.g., bar) and in the epithelium of the forming ducts (arrow). The boxed area in (C) is shown at higher magnification in (D). (E and F) P7. *Lgr5* labeling in the papilla itself appears mainly in the layer around the base of taste buds arranged in the trench walls. Fluorescence is most intense at the terminations of salivary ducts, three of which are indicated by arrowheads in (F), which is a higher magnification image of the boxed area in (E). Images represent maximal projections of 3-5 single 1 μm confocal scans. Scale bar in (A) also applies to (B) and (C).

### Expression of *Lgr5* in adult posterior papillae

*Lgr5* expression in adult animals was investigated by comparing posterior tongue sections from mouse strains harboring either the *Lgr5*-*IRES*-*lacZ* or *Lgr5*-*EGFP*-*IRES*- *CreERT2* alleles. The pattern of labeling visualized in each of these lines is very similar (Fig. 2 and Supplemental Fig. 2). The most consistent and intense labeling is seen at junctures of ducts of von Ebner’s salivary glands with the walls of the circumvallate (Fig. 2A and Supplemental Fig. 2A, B) and foliate (Fig. 2C) papillae, where it is co-extensive with the distal, excretory segments of the ducts (Fig. 2B, D and Supplemental Fig. 2A, C, F). Labeling often extends a small distance into the extragemmal taste epithelium immediately surrounding the “entry zones” where ducts merge with the papilla wall to open into the trench (Fig. 2B, D). At increasing distances from these entry zones, *Lgr5* expression, as indicated by β-galactosidase reaction product or GFP fluorescence, gradually decreases. Expression is absent from taste buds themselves.

**Figure 2.**
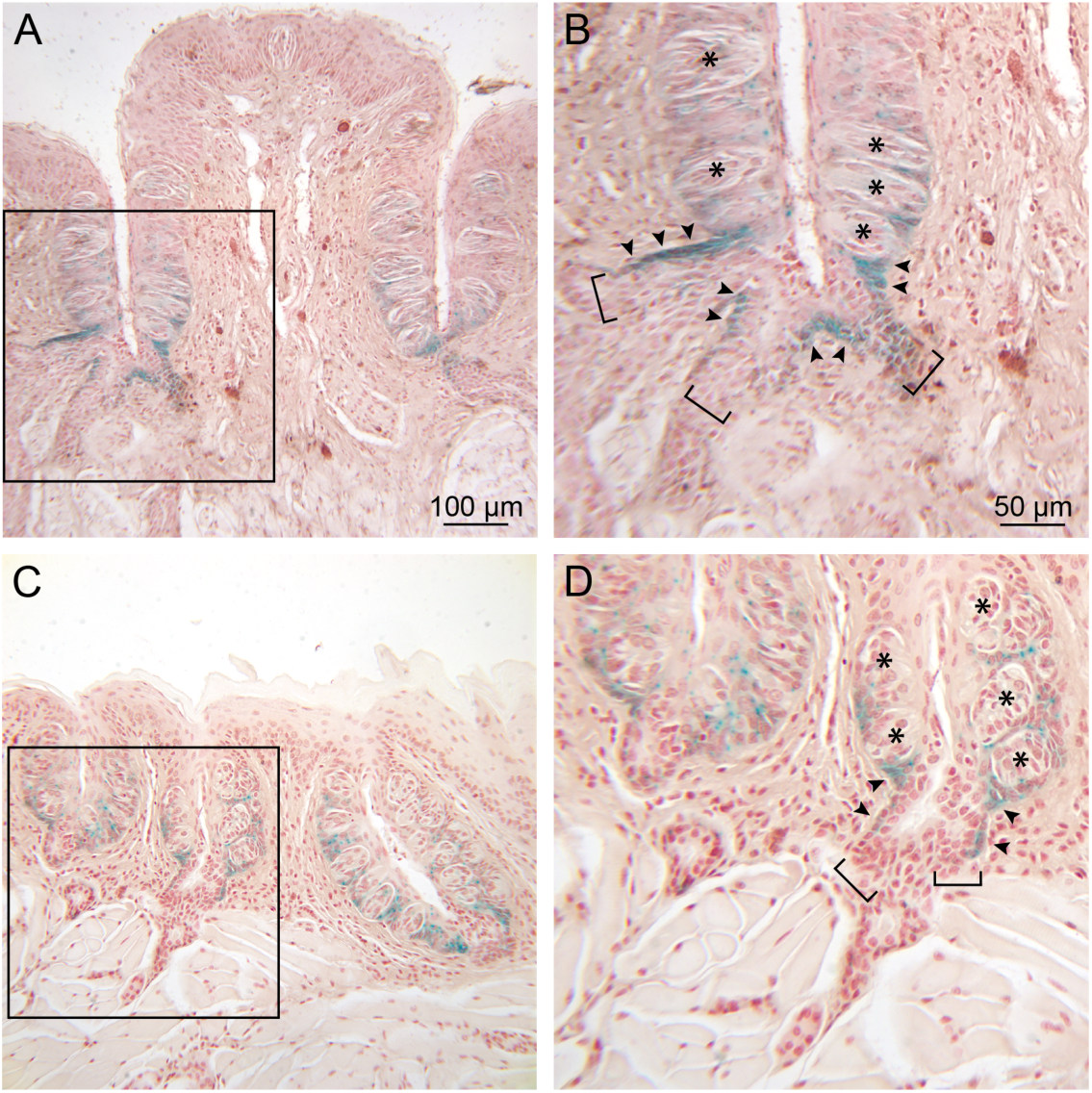
Expression of *Lgr5* in posterior tongue of adult *Lgr5*-*LacZ* reporter mice. (A and B) Circumvallate papilla. In the low power photomicrograph (A), lacZ reaction product is evident in the taste bud-containing epithelium lining the trenches and extending into extrapapillary tissue near the base of the papilla. In the boxed area, shown at 2X magnification in (B), branches of three salivary ducts (brackets) can be seen near the base of the trench. LacZ staining is intense where von Ebner’s gland ducts intersect the papilla (arrowheads) and around the base of taste buds (asterisks) in the adjacent extragemmal taste epithelium. (C and D) Foliate papillae. (C) *Lgr5* expression is shown at low power for three foliate papillae. At higher power (D), two salivary ducts (brackets) can be seen to merge as they contact the base of one papilla. *Lgr5* labeling is evident in the ducts (arrowheads) and around taste buds (asterisks), as in the circumvallate papilla in (A) and (B). Scale bars in (A) and (B) pertain to (C) and (D), respectively.

While the most prominent labeling observed in *Lgr5* reporter mice is associated with identifiable excretory ducts, it is also present in some areas near the base of taste buds when no duct is evident (Fig. 2B, D). Examination of serial sections reveals that these areas are usually found to extend from entry zones of excretory ducts that are identifiable in adjacent sections (Fig. 3E, Ei-Eviii; see description below). The number of salivary excretory ducts impinging on the papillary walls was counted in serial sections through the circumvallate papilla in five *Lgr5*-*GFP* mice. Between 14 and 18 ducts terminate at entry zones in the papilla walls in each animal. In four mice, all of the identified excretory ducts demonstrate intense *Lgr5* expression, while in the fifth animal about 20% of these ducts appear to be unlabeled (data not shown). Multiple duct entry zones are often present in the same section (e.g., Fig. 2A, B and Supplemental Fig. 2F, G). While the majority of ducts intersect the papilla wall at or near the base of the papilla, entry zones of labeled ducts can be seen distributed along medial and lateral papilla walls at loci throughout the deeper one-half of the of the papilla, but generally not more superficially. No GFP labeling appears below the two-layered excretory region in any of the papilla-directed salivary ducts (e.g., see Fig. 3A).

**Figure 3.**
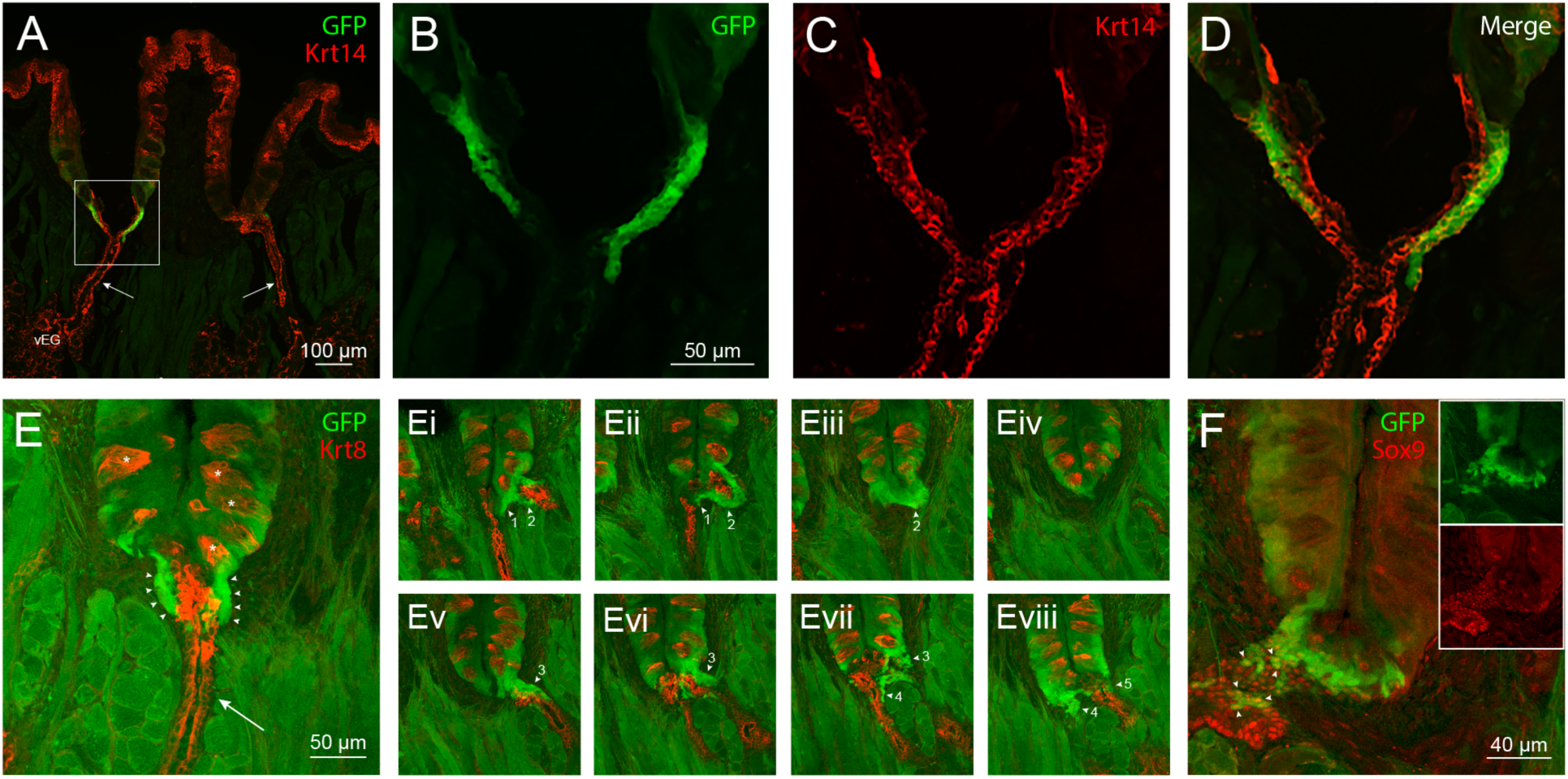
Characterization of Lgr5^+^ cells in circumvallate papillae of *Lgr5*-*GFP* reporter mice co-labeled for epithelial and ductal markers. (A-D) Immunolabeling for Krt14. In the low power confocal overlay image (A), expression of this epithelial cell marker (red) is demonstrated in von Ebner’s salivary glands (vEG), their ducts (arrows), the epithelium of the papilla and the basal layer of the surface epithelium of the tongue. A zone of high *Lgr5* expression (green) is evident in the terminal portion of the salivary duct on the left, where it contacts the base of the papilla and its lumen opens into the papilla trench. In higher magnification images of the boxed region in A, *Lgr5* expression (B) and Krt14 immunoreactivity (C) are seen to partially overlap in the distal, excretory portion of the duct (D). (E) Immunolabeling for Krt8. Anti-Krt8 reactivity (red) is located throughout the single- layered epithelium of the salivary duct (arrow) and in the internal (luminal) layer of the bi- layered excretory segment of the duct proximal to the base of the papilla. Staining for Krt8 is also seen within taste buds (asterisks). Expression of *Lgr5* (green) is limited to the external (basal) layer of the excretory segment of the duct (arrowheads) surrounding the Krt8- immunoreactive cells of the internal layer. *Lgr5* expression extends somewhat into basal portions of the adjacent papilla epithelium, but falls off in intensity with distance from the duct. (Ei-Eviii) Images of subsequent serial sections through this circumvallate papilla further illustrate the consistent association of high *Lgr5* expression with segments of at least 5 excretory ducts (numbered arrowheads) and not with the distal, single-layered portions of the ducts (red labeling only). (F) Immunolabeling for the nuclear marker Sox9. Co- localization of *Lgr5* expression (green) and Sox9 immunoreactivity (red) is seen in the excretory segment of the salivary duct where the two overlap (yellow nuclei, arrowheads). The insets show the respective single channel images for the overlay. (A-D): Single optical sections. (E and Ei-Eviii): Maximum projection of 3 optical sections. (F): Single optical sections.

### *Lgr5*-expressing cells exhibit properties of ductal cells

To better define the phenotypic properties of Lgr5^+^ cells, we compared GFP fluorescence with immunolabeling for cell type-specific markers using *Lgr5*-*GFP* reporter mice. In our experiments, keratin 14 (Krt14)-labeled cells include *Lgr5*-expressing cells of von Ebner’s gland excretory ducts, as well as non-expressing cells of gland ducts and acini and of the epithelium covering the tongue and lining papilla trenches (Fig 3A). As shown in Fig. 3B-D, Lgr5^+^ cells co-label with anti-Krt14 in the terminal region of salivary gland ducts. On the other hand, immunolabeling for keratin 8 (Krt8) localizes to taste buds and salivary gland ducts, both simple and stratified, but shows minimal overlap with *Lgr5* expression (Fig. 3Ei-Eviii). The majority of strongly *Lgr5*-expressing cells are located in a layer surrounding the single layer of Krt8-immunoreactive epithelial cells forming the excretory duct luminal wall (Fig. 3E). This labeling pattern is reproduced in multiple salivary ducts interfacing with the trench that can be observed in serial sections (Fig. 3Ei-Eviii). Expression of Sox9 (Fig. 3F), a transcription factor usually considered a marker for ductal cells, characteristically appears throughout the full extent of the salivary ducts (data not shown), notably including the *Lgr5*- expressing cells of excretory zones (Fig. 3F). Sox9 immunolabeling, however, is not seen in those cells with lower levels of *Lgr5* expression found in basal or other regions of the circumvallate papilla epithelium beyond duct entry points, or in any other taste-associated cells.

### Lgr5^+^ ductal cells are progenitors for taste bud cells

For lineage tracing studies, *Lgr5*-*EGFP*-*IRES*-*CreERT2;R26*-*mTmG* reporter mice were used. In these animals, induction of Cre recombinase following tamoxifen injection causes a switch from expression of the red fluorescent protein tdTomato to that of GFP in the cell membranes of activated cells. This allows fate mapping of cells indelibly marked by surface labeling, while at the same time monitoring continuous *Lgr5* expression, visualized as GFP targeted to the cytoplasmic compartment.

When tissues are examined at 1 day following injection, the earliest time point sampled, cells with GFP-labeled membranes are primarily *Lgr5*-expressing cells located in the excretory portion of salivary ducts (Fig. 4A, B). Induced cells are also found at the base of taste buds and in perigemmal areas (Fig. 4C, D); however, their lack of cytoplasmic GFP labeling is consistent with these being precursor cells, rather than *Lgr5*-expressing progenitor cells. By 2 days post-injection, surface-marked cells appear within taste buds (Fig. 4E). Along with perigemmal and surface epithelial cells, marked taste bud cells continue to accumulate over the next several days and, by 1 week post-injection, can be seen to comprise multiple cells in some taste buds (Fig. 4F, G). Importantly, membrane labeling persists in Lgr5^+^ excretory duct cells as late as two months following tamoxifen injection, providing evidence that this population comprises progenitor cells that are undergoing self-renewal (Supplemental Fig. 3).

**Figure 4.**
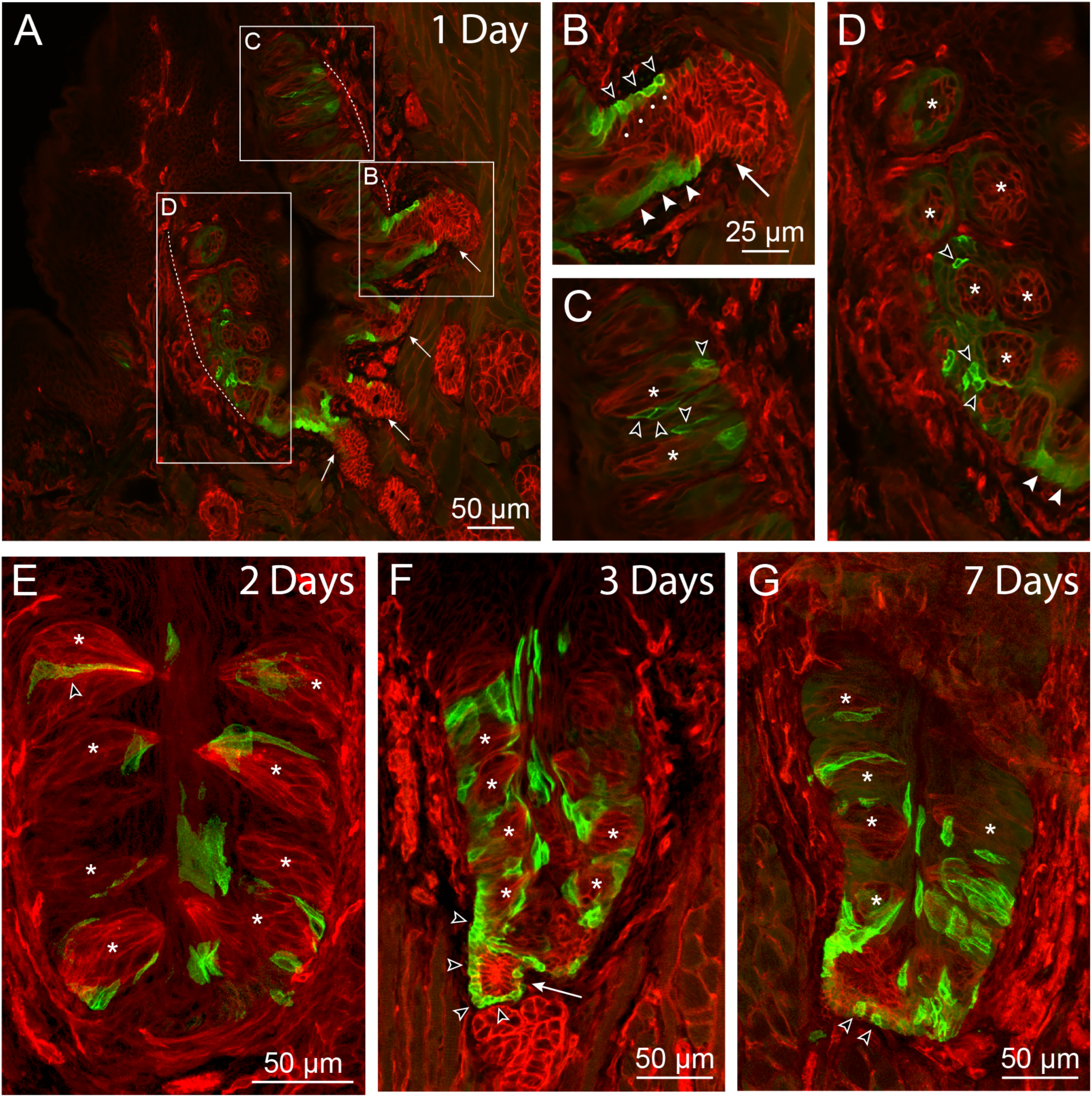
Lineage tracing of Lgr5-expressing cells in circumvallate papillae. *Lgr5*-*GFP*-*IRES*- *CreERT2;mTmG* mice were injected with a single dose of tamoxifen (day 0) and examined at the indicated time intervals for induction of Cre-mediated expression of membrane GFP. (A- D) 1 day post-injection. (A) is a low power image of one side of a papilla (dashed boundary line) with associated ducts of von Ebner’s glands (arrows). Boxed areas are seen at 2X magnification in (B), (C) and (D). Cells with induced membrane labeling are prominent at the terminations of salivary ducts, where they can be identified by the distinct circular profile (open arrowheads in B) surrounding their less intense cytoplasmic *Lgr5*-*GFP* expression. These cells, as well as uninduced Lgr5^+^ cells displaying cytoplasmic labeling only (closed arrowheads), are restricted to the basal layer of the bi-layered ductal epithelium (dots indicate luminal layer cells). Induced cells are also present in the perigemmal region in (C) and (D) (open arrowheads). In (C), asterisks indicate two longitudinally sectioned taste buds while in (D), taste buds are cut in cross section. (E) 2 days post-injection. The progeny of *GFP*-expressing cells are first seen in taste buds at this time point, illustrated by the surface- marked cell at upper left (open arrowhead). Cells with induced labeling appear at all levels within the taste epithelium. At 3 days post-injection (F), membrane-labeled cells continue to accumulate within the epithelium. A continuous line of surface-marked cells (open arrowheads) can be seen extending from the basal layer of the duct (arrow) to the basal region of the papilla below an adjacent taste bud. (G) 7 days post-injection. Duct cells with induced labeling remain (open arrowheads), with their progeny occupying taste buds as well as the surrounding papilla epithelium.

## Discussion

Collectively, our data support a role for *Lgr5*-expressing cells located in lingual salivary gland ducts as long-term progenitors/stem cells for the genesis and maintenance of posterior tongue taste buds. First, during the period of initial taste bud formation in early postnatal animals a gradual accumulation of Lgr5^+^ cells appears in nascent ducts while their numbers decrease within the papilla. Second, *Lgr5* expression persists at this location in adult animals, restricted to cells in the basal (outer) layer of terminal excretory ducts associated with von Ebner’s glands. Third, lineage tracing studies demonstrate that this specific population of cells undergoes long-term self-renewal and gives rise to differentiated cells both within and outside of taste buds in the papillae, implicating these Lgr5^+^ cells in the turnover of mature taste epithelium.

In previous expression and lineage tracing studies with *Lgr5* reporter mice, conflicting suggestions have been offered as to the location of taste stem cells. The presence of Lgr5^+^ cells surrounding the basal aspect of taste buds in adult papillae lead Takeda et al. (2013) to propose that these cells were themselves the long-term progenitors for taste cell turnover during homeostasis and in regeneration following denervation injury. On the other hand, in fate-mapping studies Yee et al. (2013) stated that “the strongest GFP signal is at the bottom of the trench area below the CV papilla and adjacent to the opening of the ducts of von Ebner’s glands”, implicating this tissue region as the source of stem cells maintaining taste bud renewal. However, in neither case did the authors identify phenotypically the specific cell type(s) expressing *Lgr5* in these varying locations. These differing observations can be reconciled by our findings that *Lgr5* labeling achieves its highest density in excretory ducts of von Ebner’s salivary glands, as well as in papillae cells directly adjoining duct entry points. Thus, the very strong *Lgr5* expression noted by Yee et al. (2013) near the trench base is consistent with the fact that this is the site where the largest number of duct entry points are found. Similarly, areas of elevated labeling adjacent to the base of taste buds elsewhere in the papilla, including in its lateral walls (Takeda et al., 2013), probably correspond to additional locations where ducts enter. Importantly, in the present study we observed that, at the earliest time point following tamoxifen injection, induced membrane labeling was apparent in the Lgr5^+^ population of cells localized to excretory ducts, as well as in extragemmal cells in immediate continuity with this ductal population. This supports the view that Lgr5^+^ cells likely move into the taste papillae at numerous locations, from progenitor niches distributed in multiple salivary ducts. Immunocytochemical studies showing that the Lgr5^+^ cells in excretory ducts co-express both Krt14 and the duct marker protein Sox9, but are devoid of Krt8, provide a signature that uniquely identifies this population of cells. It is also consistent with previous fate mapping experiments using a *Krt14-CreER* strain showing that *Krt14*-expressing cells form a lineage to taste buds (Okubo et al., 2009).

An extra-papillary source of progenitor cells capable of generating taste cells has been suggested in previous work. Using grafts of posterior tongue taste papillae into the anterior chamber of rat eyes, Zalewski (1976) found that, while taste buds disappeared in the walls of circumvallate papillae, they sometimes appeared in ductal remnants attached to the graft. The author interpreted this as suggesting a potentially important role for duct epithelium in taste bud generation. More recently, Yu et al (2020) used a *Sox10-Cre* reporter mouse strain in cell fate mapping experiments to demonstrate Sox10-Cre-labeled cells within taste buds in all three types of taste papillae. These authors found that *Sox10* transcripts were absent from the taste epithelium, but present in subepithelial connective tissue and in secretory (acinar) and duct cells of von Ebner’s glands. These authors concluded that labeled taste bud cells were, therefore, derived from precursor cells in one or more of the latter non-taste tissue compartments.

Recently, it has been possible to examine the generation of taste cells *ex vivo* in organoids from *Lgr5*-expressing cells (Ren et al., 2014; Aihara et al., 2015). Results from these experiments have shown that a self-replenishing system that generates taste cells can arise from single *Lgr5*-expressing cells isolated directly from dissociated papillae (Ren et al., 2014) or from papilla explants maintained initially in culture (Aihara et al., 2015). A consistent feature of these organoids is the expression of the progenitor/ductal cell marker Sox9 in cells of the organoid outer layer (Aihara et al., 2015). Our immunocytochemical studies have shown that the only expression of Sox9 in posterior lingual taste-associated tissues is in the highly *Lgr5*-expressing cells of the excretory duct. Furthermore, as seen in our Supplemental Fig 4, when the circumvallate papilla epidermis is stripped from the tongue using a technique similar to that used by Aihara et al. (2015) for organoid generation, the resulting preparation includes proximal segments of salivary ducts. These attached segments contain the Lgr5^+^/Sox9^+^ cells composing the excretory ducts and, thus, it is plausible that the taste cell generating functions of organoids in culture may derive from these cells.

Immunocytochemical results in the present study further characterize ductal Lgr5^+^ cells as being, in addition to Sox9^+^, also Krt14^+^. Previous lineage tracing studies using mice with an inducible *Krt14-Cre* allele have documented that taste bud cells and surrounding keratinocytes are continuously generated from long-term stem cells expressing *Krt14* in both neonatal and adult animals (Okubo et al., 2009). Based on immunolabeling with antibodies to Krt14 and other markers, these progenitors were inferred to be epithelial basal cells adjacent to taste buds. However, the Lgr5^+^/Krt14^+^ ductal cells that we describe in this study provide a possible alternative population of progenitors for the lineage of *Krt14-Cre*-labeled cells identified in taste papillae.

In conclusion, our data point to the ducts of von Ebner’s glands as a likely niche for stems cells providing renewal of taste bud and surrounding cells within circumvallate and foliate papillae. Posterior tongue taste papillae and von Ebner’s glands are closely associated with one another structurally throughout their development and, in adults, there is evidence that the two structures collaborate functionally. First, these salivary glands have been shown to secrete enzymes and binding proteins that, in addition to aiding digestive processes, might also participate in perireceptoral events influencing taste transduction (Hamosh and Burns, 1977; Roberts and Jaffee, 1986; Tremblay and Charest, 1968; Schmale et al., 1990, 1993). Furthermore, at least one brainstem autonomic reflex circuit that controls von Ebner’s gland secretory activity includes gustatory afferent input, and the potential for differential neural control of salivary secretion to modulate taste activity has also been suggested (Gurkan and Bradley, 1987). Finally, the growth factor TGFα, whose actions conceivably could support maintenance of posterior lingual taste buds, is present in von Ebner’s gland secretory vesicles (Morris-Wiman et al., 2000). Based on past studies of these interrelationships, the idea that circumvallate and foliate taste papillae and von Ebner’s salivary glands form a functional unit that could be considered a single complex organ has been proposed (Gurkan and Bradley, 1987; Sbarbati et al., 1999). Our data extend this notion by demonstrating that collaboration between these two structures is essential for taste bud turnover in posterior taste papillae and, thus, for maintaining functional integrity of this limb of the sensory system for taste.

## Acknowledgements

This research was funded in part by the National Science Foundation (DBI-REU 1062645). It was also supported by National Institutes of Health grant C06RR0306551.

The results were presented in 2013 at the 35^th^ annual meeting of the Association for Chemoreception Sciences (https://achems.org/web/downloads/programs/2013-Abstracts.pdf).

## Author Contributions

Conceived and designed the experiments: TAH DMD. Performed the experiments: TAH DMD AMD AJS. Analyzed the data: TAH DMD. Provided transgenic mice: HC JHvE. Prepared manuscript (original draft): TAH DMD. Reviewed and edited manuscript: TAH DMD AMD AJS HC JHvE.

## Supplemental Figures

Supplemental Figure 1. Visualization of *Lgr5* expression by taste papillae in neonatal and adult tongue whole-mounts from heterozygous *Lgr5*-*GFP* mice.

Supplemental Figure 2. L*g*r5 expression by circumvallate papillae of adult *Lgr5*-*GFP*-*IRES*- *CreERT2;mTmG* mice.

Supplemental Figure 3. Long-term generation of taste bud cells by Lgr5^+^ duct cells in circumvallate papillae.

Supplemental Figure 4. Circumvallate papillae isolated from mouse tongue by combined protease treatment and mechanical stripping.

