## Supplemental Figures for "Lgr5^+^ ductal cells of von Ebner’s glands are stem cells for turnover of posterior tongue taste buds"

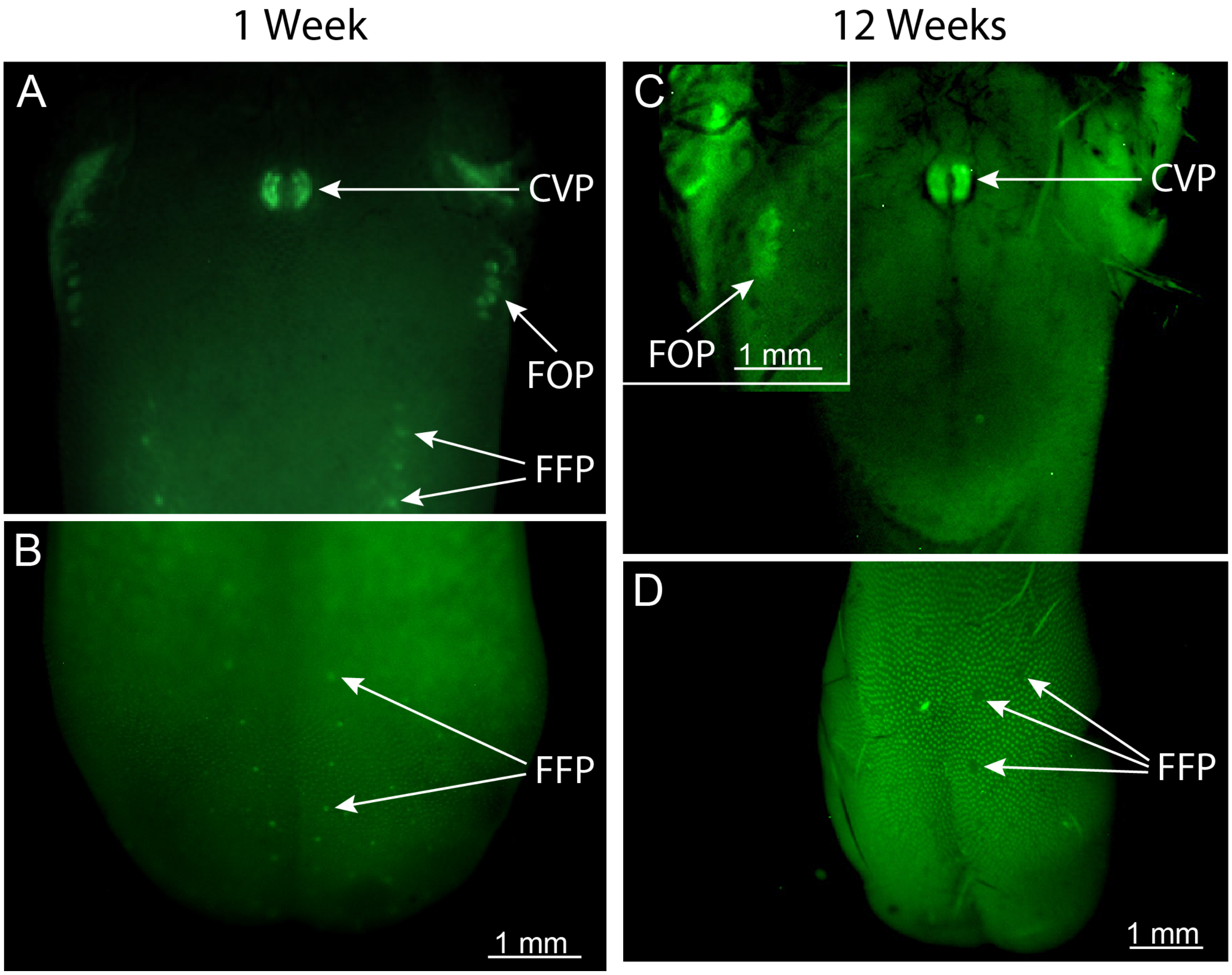

Supplemental Figure 1. Visualization of *Lgr5* expression by taste papillae in neonatal and adult tongue whole-mounts from heterozygous *Lgr5-GFP* mice. (A) and (B) 1 week. In the posterior tongue (A), GFP is localized centrally at the site of the circumvallate papilla (CVP) and laterally in the foliate papillae (FOP). Arrays of punctate label are also seen on both the posterior (A) and anterior (B) tongue at sites where single fungiform papillae (FFP) are located. (C) and (D) 12 weeks. Note that, at this time, *Lgr5* labeling is maintained in circumvallate (C) and foliate (C, inset) papillae, but is no longer evident in the fungiform papillae (D). (A) and (B) are taken from a single tongue. The main images in (C) and (D) are of the same tongue, whereas the inset image pictured in (C) is from a different mouse. The scale bar in (B) applies to (A); scale bar in (D) to (C).

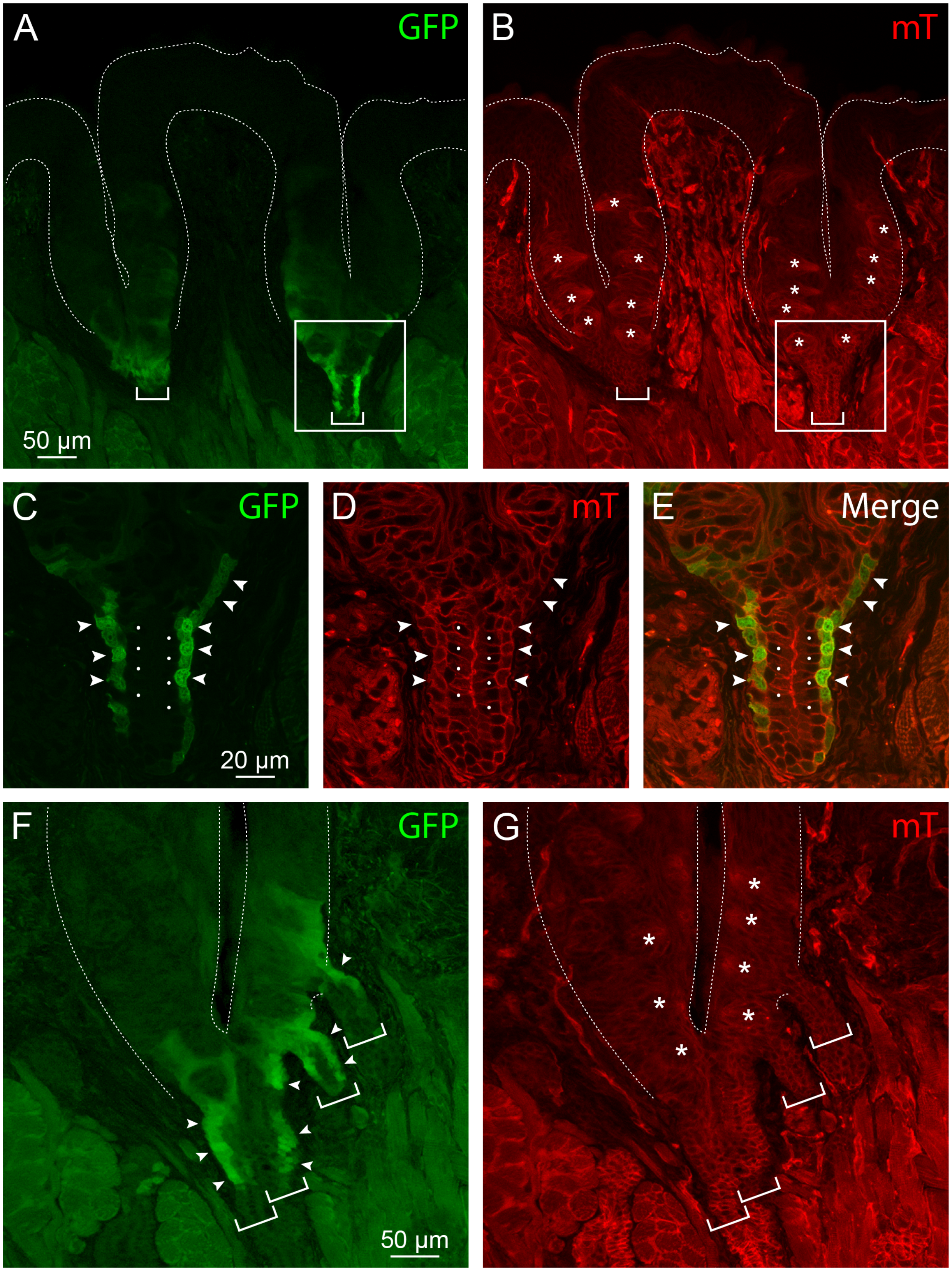
A
GFP
50 µm
B
mT
C
GFP
20 µm
D
mT
E
Merge
F
GFP
50 µm
G
mT

Supplemental Figure 2. *Lgr5* expression by circumvallate papillae of adult *Lgr5-GFP-IRES-CreERT2;R26-mTmG* mice. (A) and (B) Low power images of a single papilla. In (A), GFP fluorescence is concentrated at the base of the papilla on both sides. Cell outlines revealed by expression of membrane Tomato (mT), seen in the corresponding image (B), show that $Lgr5^+$ cells in both regions are associated with excretory ducts (brackets). The plane of section passes through the center of the duct on the right and tangentially through the epithelial wall of the duct on the left. The boxed areas in (A) and (B) are shown at 2X magnification in (C) and (D), respectively. Highly fluorescent $GFP^+$ cuboidal cells (arrowheads) occupy the outer/basal layer of von Ebner's gland excretory ducts, while columnar cells in the inner/lumenal layer of the ducts (indicated by dots) are unlabeled. (F) and (G) Intersection of excretory ducts with the circumvallate papilla. Intense GFP staining is observed in the walls of four ducts (indicated by brackets) where they merge with the papilla. While two ducts merge at the base, the others join the papilla at more superficial levels laterally. Dashed lines indicate papilla boundaries. Taste buds are indicated by asterisks. (A), (B), (F) and (G): Maximum projection of 3 optical sections. (C-E): Single optical sections.

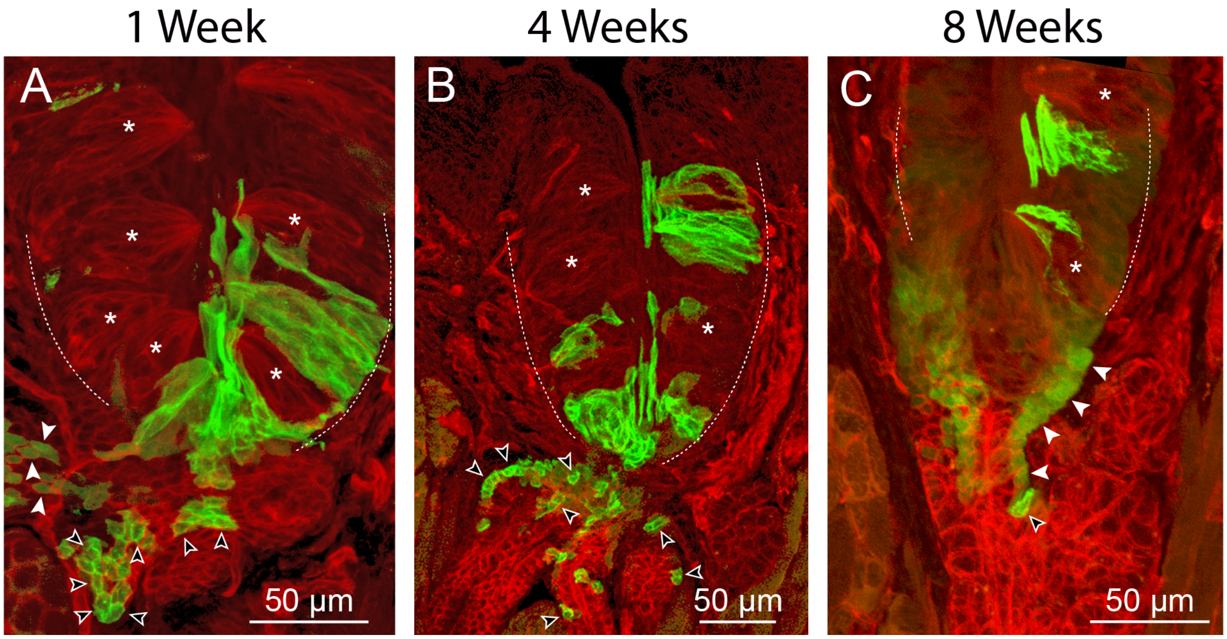
1 Week
4 Weeks
8 Weeks
A
B
C
50 µm
50 µm
50 µm

Supplemental Figure 3. Long-term generation of taste bud cells by Lgr5$^+$ duct cells in circumvallate papillae. Unlike all other reported experiments, in this case *Lgr5-GFP-IRES-CreERT2;R26-mTmG* mice were injected with two consecutive doses of tamoxifen (days 0 and 1), and then examined at the indicated times for induced membrane *GFP* expression. At 1 week (A), 4 weeks (B) and 8 weeks (C), numerous membrane-labeled cells are present in and around taste buds, including in the superficial epithelial layers lining the trench. Duct-associated Lgr5$^+$ cells, both with and without tamoxifen-induced labeling (open and filled arrowheads, respectively), are also present throughout the 8-week period, the longest time examined. Asterisks indicate individual taste buds. Dashed lines indicate the borders of the papilla.

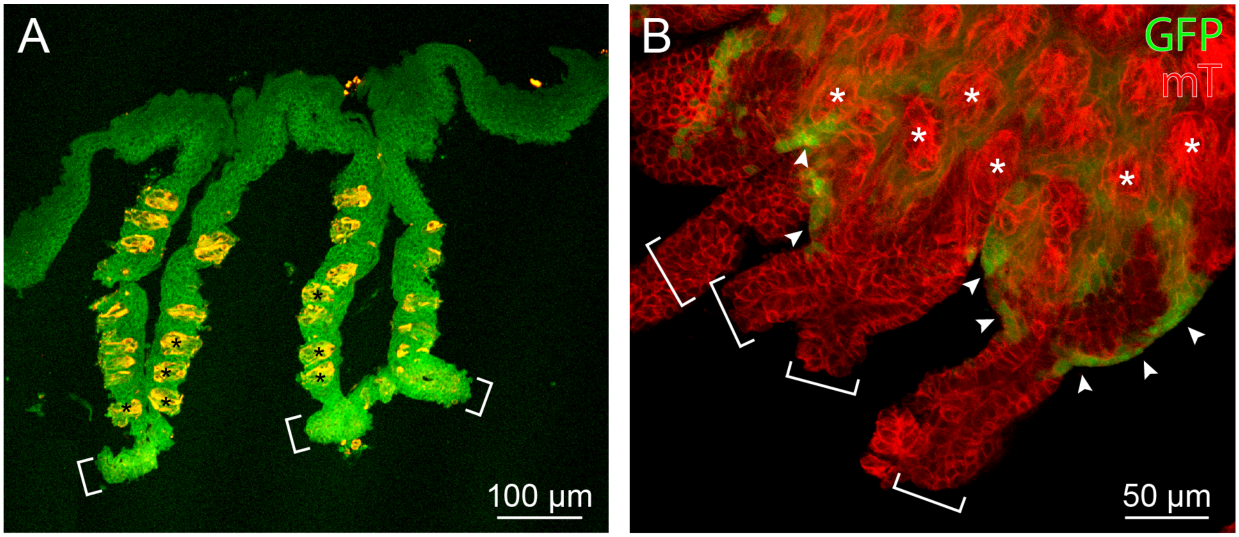

A
100 µm
B
GFP
mT
50 µm

Supplemental Figure 4. Circumvallate papillae isolated from mouse tongue by combined protease treatment and mechanical stripping. (A) Frozen tissue section from a non-transgenic animal labeled with anti-Krt8. Background tissue fluorescence (green) overlayed with Krt8 immunoreactivity (red) produces yellow-appearing taste buds (asterisks). Associated with the papilla are proximal segments of salivary ducts (brackets), which remain attached after the isolation procedure. (B) Papilla epithelium whole mount from an adult *Lgr5-GFP;R26-mTmG* mouse; all cells express membrane tomato (mT, red). $Lgr5^+$ cells (GFP, green) are present in the proximal excretory segments (arrowheads) of duct fragments (brackets) that partition with the excised tissue.
